# Ion mobility-resolved phosphoproteomics with dia-PASEF and short gradients

**DOI:** 10.1101/2022.06.02.494482

**Authors:** Denys Oliinyk, Florian Meier

## Abstract

Mass spectrometry-based phosphoproteomics has identified >150,000 post-translational phosphorylation sites in the human proteome. To disentangle their functional relevance, complex experimental designs that require increased throughput are now coming into focus. Here, we apply dia-PASEF on a trapped ion mobility (TIMS) mass spectrometer to analyze the phosphoproteome of a human cancer cell line in short liquid chromatography gradients. At low sample amounts equivalent to ∼20 ug protein digest per analysis, we quantified over 12,000 phosphopeptides including ∼8,000 class I phosphosites in one hour without a spectral library. Decreasing the gradient time to 15 min yielded virtually identical coverage of the phosphoproteome, and with 7 min gradients we still quantified about 80% of the class I sites with a median coefficient of variation <10% in quadruplicates. We attribute this in part to the increased peak capacity, which effectively compensates for the higher peptide density per time unit in shorter gradients. Our data shows a five-fold reduction in the number of co-isolated peptides with TIMS. In the most extreme case, these were positional isomers of nearby phosphosites that remained unresolved with fast chromatography. In summary, we demonstrate how key features of dia-PASEF translate to phosphoproteomics, resulting in high throughput and sensitivity.

## 1. Introduction

Protein phosphorylation is one of the most prevalent and most studied post-translational modifications (PTMs)^1^. The reversible addition of a phosphoryl group to amino acid chains alters the physicochemical properties of a protein and, in turn, modulates its localization, interactions or enzymatic activity. Deregulated signal transduction via phosphorylation has been linked to numerous diseases and is one of the hallmarks of cancer^2-5^. Advances in mass spectrometry (MS)-based proteomics, including the development of efficient protocols for phosphopeptide enrichment and sophisticated computational algorithms for the identification, quantification and localization of modification sites resulted in reports of >150,000 distinct phosphorylation sites on >75% of the human proteome^6-9^. However, for the vast majority, their cellular function and underlying kinase-substrate networks remain elusive^10^.

To disentangle the functional role of specific phosphosites, there is a trend towards experimental designs that investigate the dynamic regulation of the phosphoproteome in many different cellular states, compartments and conditions^11-13^. This spurs ongoing efforts to increase the throughput of phosphoproteomics experiments, for example by means of shortening liquid chromatography gradients^6^. However, shorter gradients entail increasing signal complexity per time unit, and often decreasing quantitative accuracy. Also, reproducibility can be compromised in classical data-dependent acquisition modes as the increasing number of co-eluting peptides exceeds the sequencing speed of the mass spectrometer. In part, this is addressed by the advent of data-independent acquisition (DIA) strategies that fragment multiple precursor ions in parallel to trace peptide fragment ions along their chromatographic elution peak. This is typically achieved by cycling the quadrupole mass filter through a pre-defined list of precursor windows, collectively covering the full mass range of interest^14^.While narrower isolation windows facilitate peptide identification, an immediate consequence of the acquisition scheme is that a peptide of interest is sampled in only every *n*-th scan.This also applies to more recently developed methods in which the quadrupole is scanned continuously rather than in discrete steps to enable extremely fast acquisition cycles compatible with chromatographic gradients of five minutes or less^25, 26^.In contrast, we have recently demonstrated that combining trapped ion mobility spectrometry (TIMS) and parallel accumulation – serial fragmentation (PASEF) with DIA in a novel scan mode termed dia-PASEF can dramatically increase the utilization of the ion beam^15,16^. This is because in dia-PASEF, all precursor ions are trapped in the TIMS device in parallel and sequentially released as a function of their ion mobility. Due to the correlation of peptide mass and mobility, the precursor selection can be programmed to follow the population of peptide ions as they exit the TIMS device towards the downstream quadrupole time-of-flight (QTOF) mass analyzer.

Incorporating ion mobility separation into conventional LC-MS setups also adds an extra dimension of separation by size and shape in the gas phase^17-18^. We and others have estimated that, in proteomics practice, TIMS increases the analytical peak capacity about ten-fold^19,20,34^. Intriguingly, phosphorylation can have a distinct influence on the gas phase conformation of a peptide as measured by its collisional cross section (CCS)^21^. Analysis of a large number of phosphopeptides revealed that their distribution in the ion mobility vs *m/z* space is shifted towards more compact conformations as compared with unmodified peptides, suggesting sequence-specific interactions of the phosphoryl group with basic amino acids^22-23^.

Here we reasoned that the increased ion utilization of dia-PASEF combined with the ion mobility resolution of TIMS should facilitate high-throughput phosphoproteomics. To this end, we analyze the phosphoproteome of a human cancer cell line with gradients from 7 to 60 minutes and assess the separation of phosphopeptides in both dimensions, liquid chromatography and trapped ion mobility.

## 2. Materials and methods

### Human cell culture

Human epithelial cervix carcinoma HeLa cells were cultured in RPMI 1640 medium supplemented by 10% fetal bovine serum, 2% penicillin/streptomycin and 5 µl/ml plasmocin at 37 °C in an incubator with 5% CO_2_. Cells were harvested at approximately 80% confluency by incubation with 0.25% trypsin/EDTA, transferred to 15 mL falcon tubes, washed twice with TBS, pelleted by centrifugation for 5 min at 1000 g and stored at −80 °C until further use.

### Cell lysis and protein digestion

We lysed the cell pellets by adding SDC lysis buffer (4% sodium deoxycholate, 100mM Tris-HCl pH 8.5) and boiling for 10 min at 95 °C, before high-energy sonication to shear DNA (Diagenode Bioruptor pico). The protein concentration was determined through a BCA assay (Thermo Fischer). Protein disulfide bonds were reduced by adding tris(2-carboxyethyl)phosphine and the free cysteine residues were carbamidomethylated with 2-chloroacetamide at final concentrations of 10mM and 40mM, respectively, for 5 min at 45 °C. Finally, lysates were digested overnight with trypsin and Lys-C at an enzyme/protein ratio of 1:100 (w/w) for both.

### Phosphopeptide enrichment

We started from 200 µg of proteome digest to enrich phosphorylated peptides according to the ‘EasyPhos’ protocol as described^9^. In short, after mixing the peptides with isopropanol (ISO) and EP enrichment buffer (48% TFA, 8mM KH2PO4), phosphopeptides were enriched within 5 min at 1500 rpm and 40 °C using 5 mg of TiO_2_ beads at a concentration of 1 mg/µL in 6% TFA/80% ACN (vol/vol). To remove unspecific binders, we washed the beads with 5 mL of 5% TFA/60% ISO (vol/vol). The phosphopeptides were then eluted with the EP elution buffer (40% ACN, 15% NH4OH) and vacuum-concentrated in a SpeedVac for 30 min at 45 °C. Subsequent desalting and peptide purification was performed on SDB-RPS StageTips. For LC-MS analysis, the dried samples were reconstituted in MS loading buffer (0.1% TFA/2%ACN (vol/vol)) to a final concentration equivalent to 20 µg/µl protein input amount.

### Liquid chromatography-MS analysis

Nanoflow reversed-phase liquid chromatography was performed on a nanoElute system (Bruker Daltonics). Peptides were separated within 7, 10, 15, 21, 30, 60 min gradients at flow rates of 1.5, 1, 0.8, 0.8, 0.8, 0.5 µL/min, respectively, on a 8 cm x 75 µm column with 1.9 µm ReproSil-Pur C18 - AQ particles (PepSep). Mobile phases A and B were water with 0.1 vol% formic acid and ACN with 0.1 vol% formic acid, respectively. The LC system was connected online to a hybrid TIMS QTOF mass spectrometer (Bruker timsTOF Pro) via a CaptiveSpray nano-electrospray source. The mass spectrometer was operated in dia-PASEF mode, for which we defined isolation windows in the *m/z* vs. ion mobility plane adjusted to expected precursor ion density for phosphopeptides using the pydiAID software^37^. We sampled an ion mobility range from 1/K_0_ = 1.43 to 0.6 Vs cm^-2^ using equal ion accumulation time and ramp times in the dual TIMS analyzer of 100 ms each. The collision energy was lowered as a function of increasing ion mobility from 59 eV at 1/K_0_ = 1.4 Vs cm^-2^ to 20 eV at 1/K_0_ = 0.6 Vs cm^-2^. For all experiments, we calibrated the ion mobility dimension linearly using at least two out of three ions from Agilent ESI LC/MS tuning mix (*m/z*, 1/K_0_: 622.0289, 0.9848 Vs cm^-2^; 922.0097, 1.1895 Vs cm^-2^; and 1221.9906, 1.3820 Vs cm^-2^).

### Raw data processing

The dia-PASEF raw files were processed in Spectronaut v15.1.0.27012 in library-free mode (‘directDIA’) using the default settings, including a 1% false discovery rate at precursor and protein levels. We defined Carbamidomethylation (C) as a fixed modification and Acetyl (Protein N-term), Oxidation (M) and Phospho (STY) were set as variable modifications. For the database search, we used the UniProt human proteome reference FASTA file downloaded on 15th of September 2021. To analyze phosphopeptides, we activated the PTM localisation mode and defined a threshold of 0 for all phosphopeptides, 0.75 for class I phosphopeptides and 0.99 for the analysis of phosphopeptide positional isomers. Data filtering for quantification was set to Q-value, normalization to Automatic and Cross Run Normalization was enabled. To report only unique phosphosites, we collapsed duplicate entries from proteins within the same protein group in the Spectronaut PTM site output table.

To estimate spectral complexity on the MS1 level, we processed one raw file per gradient with MaxQuant v2.0.3.0 to perform feature detection. We then analyzed the *allPeptides*.*txt* file and iterated through all scan indices to extract the number of detected features per scan. To estimate the precursor density per dia-PASEF isolation window, we *(i)*filtered the detected features by *m/z* as defined in the dia-PASEF acquisition scheme(‘without ion mobility’) and *(ii)*calculated for each *m/z* window the average number of features with overlapping ion mobility peak boundaries at full width at half maximum (‘with ion mobility’).

### Bioinformatics analysis

Data analysis and visualization was performed using custom scripts in R (4.0.1) and Python (3.8.8) with packages data.table (1.14.2), dplyr (1.0.7), ggplot2 (3.3.5), tidyR (1.1.14), patchwork (1.1.1), pandas (1.1.5), numpy (1.22.2), plotly (5.4.0), scipy (1.7.3).

## 3. Results& Discussion

### Workflow for high-throughput phosphoproteomics

Scaling the throughput of phosphoproteomics experiments involves shortening the analysis time per sample as well as a reduction of starting material to facilitate parallel sample processing and limit costs. To assess the performance of dia-PASEF in such a scenario, we enriched phosphopeptides from human cervix carcinoma cells and aliquoted them into equivalents of 20 µg starting material (the protein mass of∼100,000 cells). This approach mimics 10 to 50 times less sample than typically recommended in state-of-the-art enrichment protocols, while reducing confounding factors in our analytical assessment of dia-PASEF^6, 9, 27^. Further, to explore the limits of sample throughput, we analyzed the enriched phosphopeptides within 7 to 60 minutes LC-MS time (**Fig. 1**).We designed the LC gradients to distribute the ion current evenly within the available separation time, while minimizing void times and sample-to-sample carry-over with a short 8 cm column. Including overhead times, for example due to autosampler operations and column equilibration, this setup allows analyzing between 20 and 120 samples per day with nanoflow-sensitivity.

**Figure 1.**
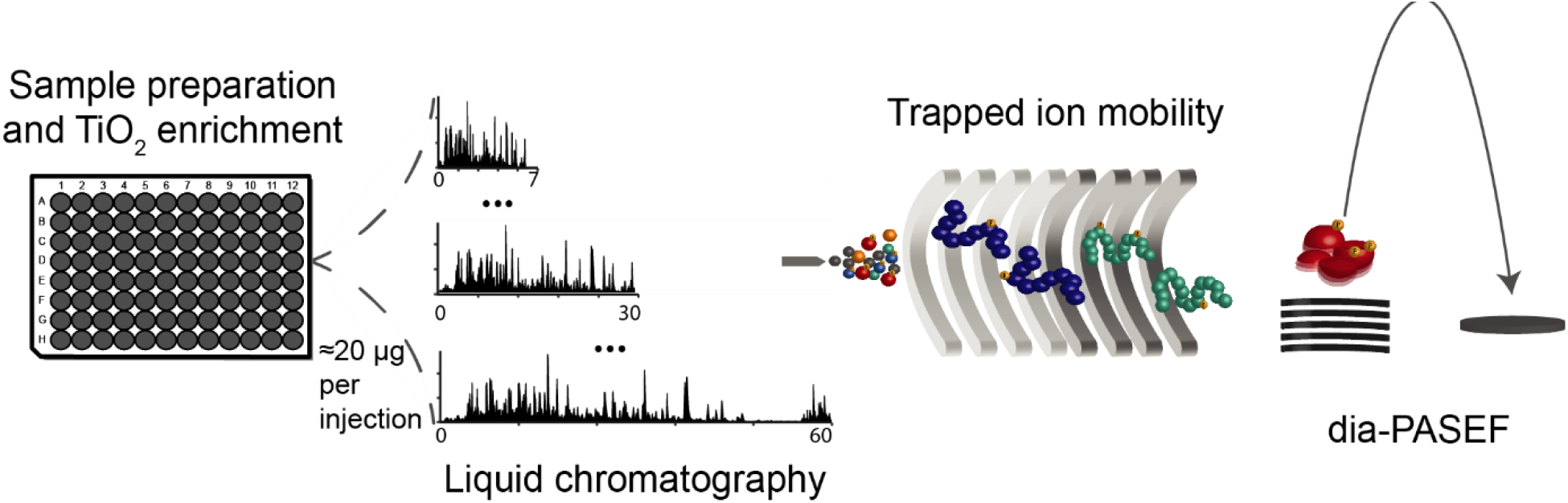
Schematic workflow to assess the performance of dia-PASEF for high-throughput and high-sensitivtiy phosphoproteomics. Equal aliquots of phosphopeptides enriched from a HeLa cell lysate are separated in different LC gradients from 7 to 60 min and analyzed with a trapped ion mobility – quadrupole time-of-flight mass spectrometer using the dia-PASEF acquisition mode.

Steeper gradients compress the peptide signal into narrower elution peaks. On the one hand, this is favorable because it increases the signal intensity. One the other hand, it requires faster MS methods to accurately reconstruct the peak shape for quantification. A particular feature of dia-PASEF is that acquisition schemes can be readily tuned for high throughput applications without reducing selectivity^15^. This is because ion mobility separated precursors from multiple DIA windows are fragmented sequentially within a single PASEF scan. Here, we employed a dia-PASEF acquisition scheme with a cycle time of ∼1.4 s comprising one TIMS-MS survey scan and 24 isolation windows allocated to 12 dia-PASEF scans. To account for well-established differences in the gas phase conformation of phosphorylated and unmodified peptides, we positioned the isolation windows in the *m/z* vs. *1/K*_*0*_ space in a way that they optimally cover the area occupied by phosphopeptides and adapted the window width to the local peak density (Skowronek *et al*., in preparation).

To process the dia-PASEF raw data files we chose a library-free approach in which pseudo-MS/MS spectra are built from correlating precursor and fragment ions, and matched to an *in silico* digest of the reference proteome^35^. In our hands, this streamlines the overall workflow and guarantees flexibility as it avoids the generation of project-specific spectral libraries. Of note, this approach holds particular promise for phosphoproteomics as it can detect rare or low-abundance phosphosites that might be absent from an experimental library^6^.

### Phosphoproteome coverage in short gradients

Having outlined a high-throughput dia-PASEF strategy for phosphoproteomics, we first evaluated its performance in terms of phosphopeptide identifications for increasingly fast separations. To set a reference, we analyzed the HeLa phosphoproteome with a 60 min gradient (∼20 samples per day).We identified in total∼12,000 phosphopeptides from quadruplicate injections of the equivalent of 20 µg protein digest. Out of these, the Spectronaut software localized ∼8,000 with a probability >0.75 (class I sites) at a specific serine, threonine or tyrosine residue in one of 2,630 proteins (**Fig. 2A**). The7 min gradient still resulted in∼9,200 phosphopeptides and 2,349 phosphoproteins, however, at a 6-fold increased sample throughput (**Fig. 2B**).Most notably, this covered the major signaling pathways, indicating that at a throughput of 120 samples per day we achieved a phosphoproteomic depth that can already provide relevant biological insight (**Suppl. Fig. 1**).

**Figure 2.**
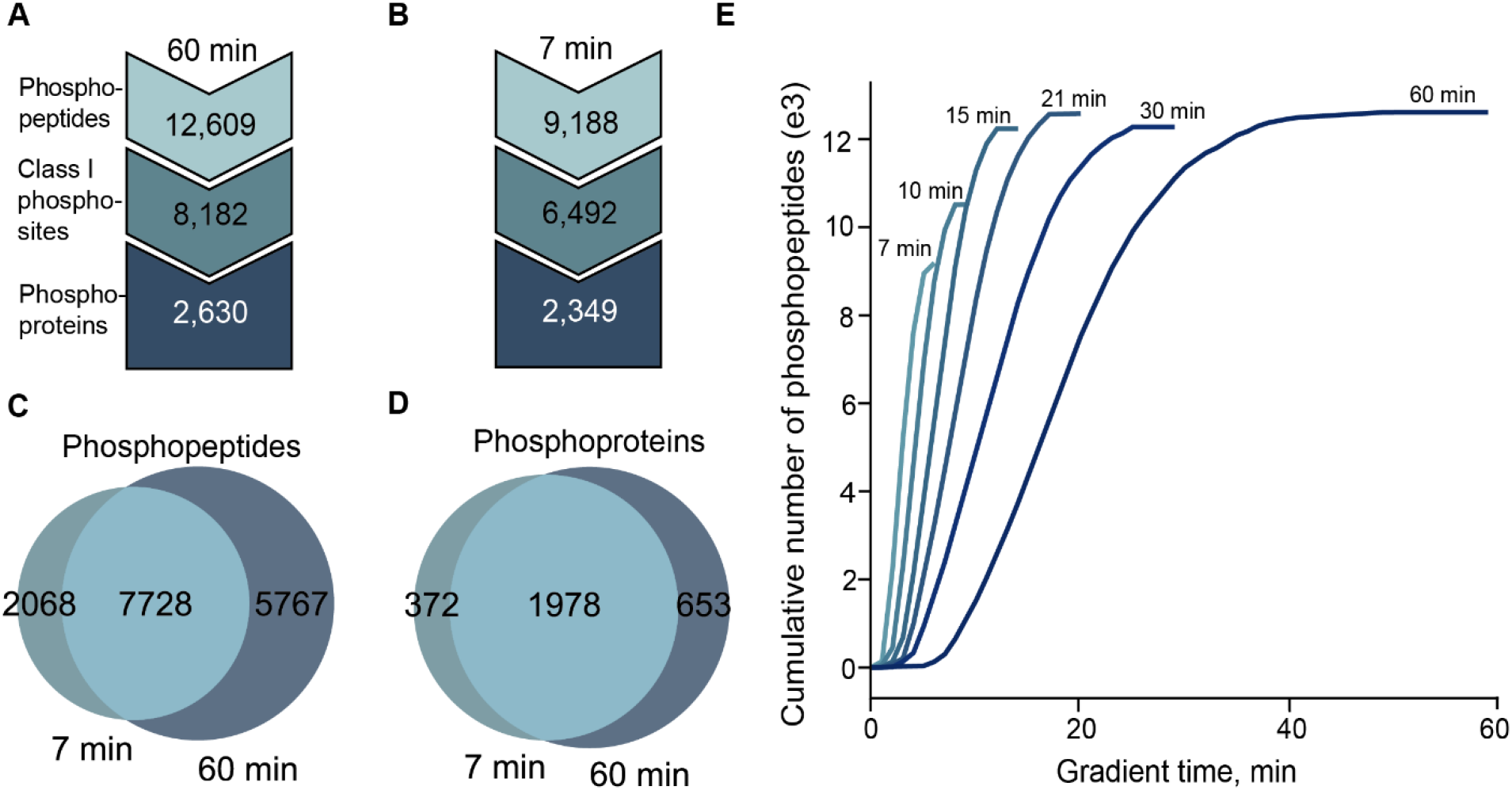
dia-PASEF phosphoproteomics with fast chromatographic separation. **A** Total number of identified phosphopeptides and phosphorylated proteins in quadrupulicate analysis with 7 min LC gradients, injecting aliquots equivalent to 20 µg protein extract. **B** Same as A, but 60 min LC gradients. **C** Overlap of phosphopeptides between 7 and 60 min LC gradients. **D** Overlap of phosphorylated proteins between 7 and 60 min LC gradients. **E** Cumulative number of identified phosphopeptides as a function of gradient time.

Proteome analysis typically aggregates multiple peptides per protein. This redundancy is missing in phosphoproteomics as each peptide typically represents an individual modification site. For this reason, we next assessed the reproducibility on the peptide level. For both gradients we found virtually complete data matrices (>99%) and the vast majority of phosphopeptides were detected in all four replicates. We trace this result back to the combination of consistent data acquisition with dia-PASEF and highly specific data analysis with the library-free approach. Reassuringly, we also found a high overlap between the two gradients (**Fig. 2C, D**). On both the peptide and protein level, about 84% of the identifications in the 7 min gradient were in common with the 60 min gradient, while peptides uniquely identified in only one of the gradients showed a bias towards lower abundance (**Suppl. Fig. 2A, B**).

One of the key features of DIA is parallel peptide sequencing. To explore the current limits thereof, we plotted the cumulative number of identified phosphopeptides as a function of the gradient time (**Fig. 2E**). For the 60 min gradient, our analysis showed an identification rate of ∼300 peptides/min in the center of the gradient and revealed a slight identification bias towards the hydrophilic part of the gradient. Reducing the gradient time to 30 min resulted in essentially the same number of identified peptides, however, utilizing the MS sequencing capacity more efficiently as evident from a steeper curve of cumulative identifications. Further compressing the peptide elution window in 21 and 15 min gradients, we achieved sequencing rates of 1,200 and 1,400 peptides/min, still with no visible loss in phosphoproteome coverage. This highlights the advantages of dia-PASEF for short gradients and indicates diminishing returns for longer gradients, in particular for low sample amounts and lower-complexity samples. Even shorter gradients of 10 and 7 min increased the sequencing rate up to 1,800 peptides/min for the 7 min gradient, which, however, was no longer sufficient to fully compensate for the increasing sample complexity. This suggests that we reached a point at which separation power becomes a limiting factor.

### Ion mobility separation of co-eluting and co-isolated peptides

Resolving peptides of similar mass and retention time is not only important for identification, but also indispensable for accurate quantification. This prompted us to investigate the analytical peak capacity in more detail. In the 60 min gradient, chromatographic peak widths were on average 11 seconds, which equates a theoretical capacity of ∼330 peaks (**Fig. 3A**). Decreasing the gradient time coincided with gradually narrower peptide elution peaks. For this reason, the peak capacity of the 30 min gradient was about two-thirds of the 60 min gradient and the peak capacity of the 7 min gradient was ∼90 with an average peak width of 5 s. In other words, the complexity of the sample as seen by the mass spectrometer was theoretically up to 3-fold higher in the 7 min gradient as compared with the 60 min gradient. To this end, we used the MaxQuant feature detection algorithm to calculate the actual number of isotope patterns detected in every TIMS-MS survey scan as a proxy for spectral complexity (**Fig. 3B**). This peaked around 1,000 per spectrum for the 60 min gradient, while shorter gradients increased the peptide density up to 1,600 detectable features per spectrum in the 7 min gradient.

**Figure 3.**
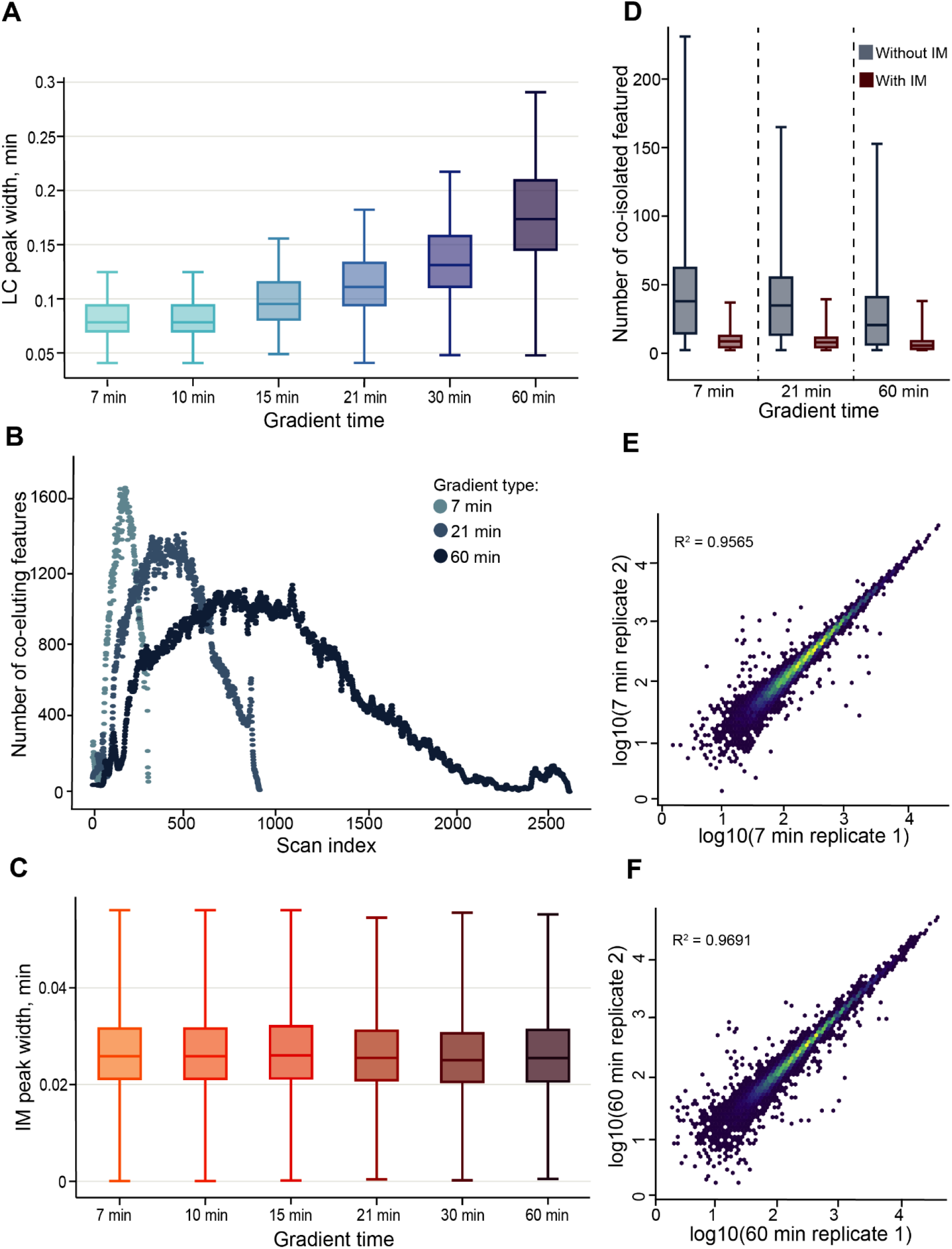
Accurate quantification of the phosphoproteome with dia-PASEF. **A** Boxplots ofchromatographic peak widths at 5% maximum intensity of all identified phosphopeptides insix different LC gradients. **B** Number of co-eluting features per TIMS scan in 7, 21 and 60 min gradients. **C**Ion mobility peak widths at 5% peak intensity of all identified phosphopeptides. **D** Number of co-isolated features per dia-PASEF isolation window with and without ion mobility separation for 7, 21 and 60 min gradients. **E**Measured phosphopeptide intensities intwo replicate injections with the 60 min gradient. **F**Same as E, but 7 min gradient.

Next, we asked to what degree the additional ion mobility separation mitigates the lower resolving power of shorter gradients. TIMS is performed post-column in the gas phase and should therefore be independent from chromatographic parameters. Indeed, we observed a very consistent ion mobility peak width distribution across all experiments, corresponding to a TIMS peak capacity of 32 (**Fig. 3C**). That is, while less peptides can be resolved from each other chromatographically, they are still effectively separated in the ion mobility dimension. As a ballpark figure, correcting for the correlation of peptide mass and mobility, we estimate that the two-dimensional peak capacity of the 7 min gradient and 100 ms TIMS scans is ∼800, which is two to three times higher than the 60-min chromatographic separation alone.

To see how these considerations translate to dia-PASEF experiments, we extracted the number of features contained in each isolation window for the 7, 21 and 60 min gradients, with and without considering the ion mobility separation (**Fig. 3D, Methods**). In line with the above, we observed an inverse relationship between gradient length and the number of features per isolation window. This is to the extent that we found cases of over 200 species overlapping in a single *m/z* window of the 7 min gradient. Conversely, considering only precursors with overlapping ion mobility peaks (FWHM)in this analysis reduced the average number of co-isolated features consistently by a factor of five for all gradients. We thus conclude that the additional TIMS dimension can indeed resolve interfering signals that would otherwise impede peptide identification and quantification. Confirming this result, we found an excellent quantitative precision for the 60 min gradient (median peptide CV 8.5%) as well as for the 7 min gradient (median CV 8.9%), covering a similar abundance range (**Fig. 3E, F**).

### Resolving isobaric positional isomers

A particular attraction of ion mobility in phosphoproteomics is the possible separation of isobaric positional isomers,as these have the same mass, typically elute in close proximity and share many fragment ions^29^.We therefore investigated whether dia-PASEF is successful in discriminating positional isomers even though we deliberately lowered the chromatographic resolution to increase throughput and only analyzed low sample amounts. To obtain a coherent and sufficiently large subset of phosphopeptides for this analysis, we focused on pairs of peptides with one phosphoryl group on alternative serine or threonine sites within the same sequence. Applying a stringent localization probability cut-off >0.99, we were left with 2,771 high-confidence isobaric pairs matched by charge state and across all gradients, corresponding to 657 unique modified sequences. For each of these, we calculated pairwise resolution values in both LC and TIMS dimensions (**Fig. 4A**). Chromatography baseline resolved 2,394 positional isomer pairs (86%), and the median chromatographic resolution was4.6. Gradients of 15 min and longer accounted for the largest proportion (412 to 442 resolved isomers), while the 7 min gradient was still sufficient to resolve 313 isomers (**Fig. 4B**).

**Figure 4.**
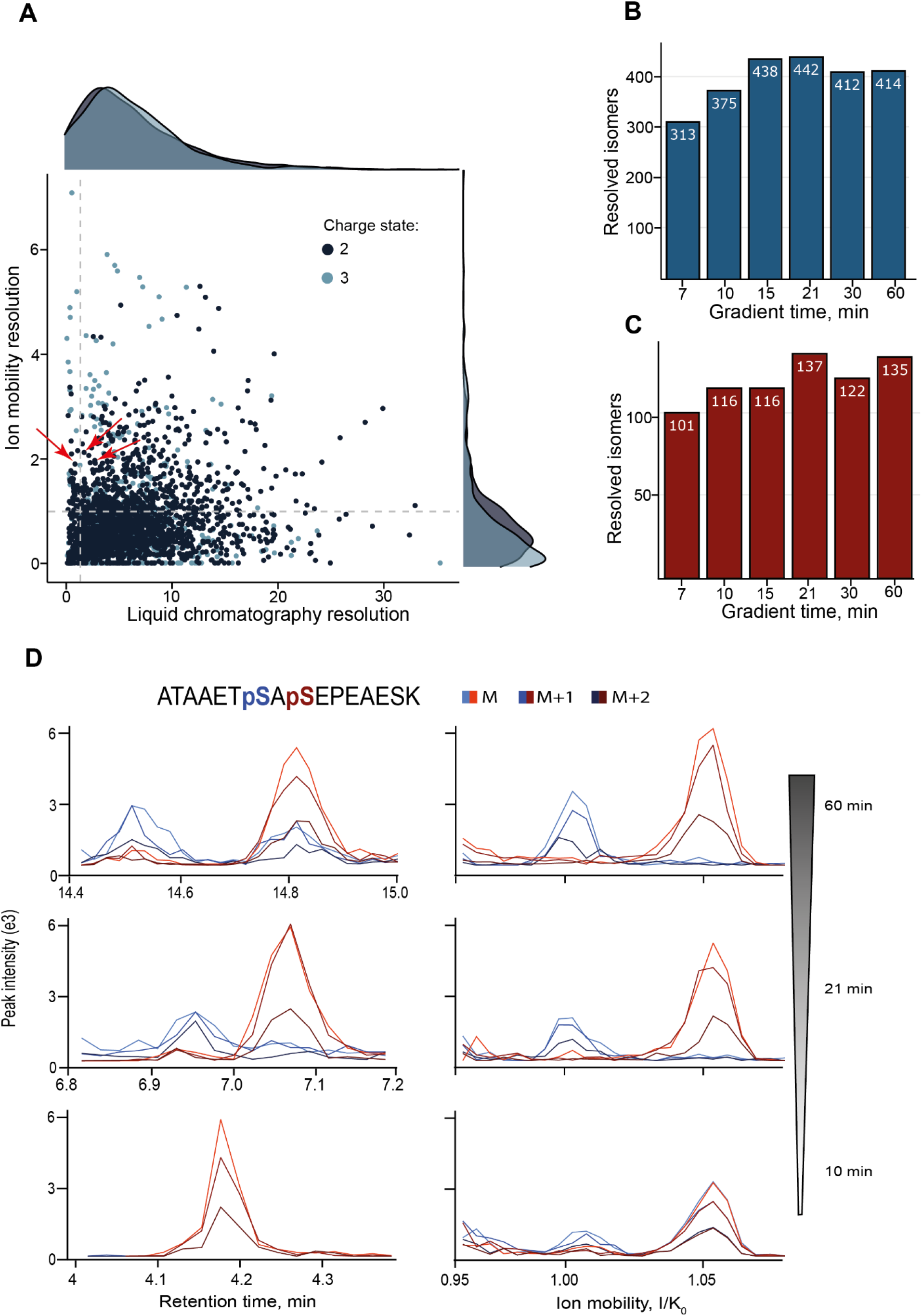
Two-dimensional separation of isobaric phosphopeptides. **A** Pairwise comparison of positional isomer separation in liquid chromatography and ion mobility (n = 2771 pairs).**B** Baseline resolved pairs of positional isomers by chromatography with different gradients. **C** Same as B, but resolved by ion mobility. **D** Extracted ion chromatograms and mobilograms for an example of isobaric positional isomers separated in progressively shorter gradients of 60, 21 and 10 min and 100 ms TIMS scans. Red arrows in A indicate the position of the peptide pair shown in D.

Despite its compact design, TIMS can be readily tuned to a high ion mobility resolution by adjusting the scan rate^36^. In our case, 100 ms scans baseline-resolved in total 727 pairs of isobaric positional isomers baseline across all gradients (median resolution 0.6), which is about one quarter of all pairs (**Fig. 4A**).We noticed a somewhat higher resolution for triply charged than for doubly charged peptides, likely corresponding to an increased conformational flexibility of longer phosphopeptides in the gas phase. Similar to the overall trend in identifications, the number of ion mobility-resolved positional isomers ranged from 101 for the 7-min gradient to 137 for the 60-min gradient (**Fig. 4C**).Of note, among them were 36 pairs of isobaric phosphopeptides that were not resolved at all by chromatography.

To illustrate this further, we extracted ion chromatograms for a representative pair of isobaric positional isomers (ATAAETpSASEPEAESK and ATAAETSApSEPEAESK, red arrow in **Fig. 4A**) that was identified consistently across the 10, 21 and 60 min gradients with a localization probability >0.99 (**Fig. 4D, Suppl. Fig. 3**). In the 60 min gradient, both isomers were baseline separated and eluted about 20 seconds apart. Likewise, the corresponding ion mobility spectra revealed a ∼5% difference in *1/K*_*0*_ indicating that the two isomers differ substantially in their size and shape in the gas phase. With decreasing chromatographic resolution, the elution peaks started to collapse and finally overlapped in the 10 min gradient. At the same time, the ion mobility resolution remained constant, allowing us to distinguish both isobaric isomers regardless of their chromatographic separation. Furthermore, their relative abundance was similar in all gradients, adding further evidence to our notion that the quantitative accuracy is preserved in dia-PASEF even for short gradients.

## 4. Concluding remarks

Modern MS-based phosphoproteomics measures tens of thousands of protein phosphorylation events in biological samples. However, accurate and consistent quantification remains challenging for large sample cohorts and limited sample amounts. To address this, we here developed a workflow that leverages the inherent speed and sensitivity of dia-PASEF to analyze cellular phosphoproteomes from tens of µg of protein material in short gradients without the need for a project-specific spectrum library.

A recent study by Bekker-Jensen *et al*. highlighted advantages of data-independent acquisition for rapid phosphoproteome profiling with the Orbitrap mass analyzer^6^. We reasoned that faster time-of-flight mass spectrometers should accentuate these advantages even more because, in principle, they can achieve very short cycle times that are sufficient to accurately quantify chromatographic peaks of just a few seconds with high mass resolution^25^. In classic DIA, however, faster scan cycles come at the expense of selectivity (wider isolation windows) or sensitivity (shorter scan time)^14,30^.In contrast, by synchronizing precursor selection and ion mobility separation, dia-PASEF increases the ion sampling efficiency without compromising selectivity^15^. In this study, we demonstrated that these advantages translate directly to phosphoproteomics. Highly parallelized sequencing of over 1,400 phosphopeptides per minute allowed us to reduce the analysis time 4-fold at no loss in coverage. At the same time, we found that TIMS effectively reduces the overlap of co-eluting phosphopeptides in the same precursor window, which is in line with similar simulations for unmodified peptides^19,20^.This can facilitate fragment ion assignment and resolve interferences for accurate quantification, which is of particular importance for short gradients. Comparable to Bekker-Jensen et al., we quantified about 12,000 phosphopeptides in quadruplicate 15-min gradients, however, injecting five to ten times less material per analysis, and achieving a virtually complete phosphopeptide data matrix. Library-based data analysis could increase the phosphoproteome coverage further, yet at a potentially lower level of data completeness and with increased experimental complexity and cost. We also note that the sample throughput could be readily increased while maintaining the active gradient time using tailored liquid chromatography systems^28^. Likewise, our workflow should benefit immediately from the development of ultra high-sensitivity timsTOF systems^31^.

In principle, DIA isvery well suited to resolve individual phosphorylation sites because the mass spectrometer records chromatographic profiles of all detectable fragment ions. This is countered by the sheer complexity of DIA spectra, which turns the precise assignment of site-specific fragment ions into a difficult task^6,29,32^. As shown here, increasing throughput with shorter gradients can exacerbate this situation. Nevertheless, our workflow gave rise to over 6,000confidently localized phosphosites even with 7-min gradients. Analysis of a subset of isobaric positional phosphopeptide isomers highlighted that LC and TIMS can be complementary, opening up the possibility of increasing chromatographic throughput and at least to some extent still being able to resolve isomeric peptides. It could also be interesting to inspect the separation of positional isomers in the ion mobility dimension with dedicated software packages such as Thesaurus^29^, however, these do currently not support dia-PASEF data.

In conclusion, our study provides a path towards high throughput and high sensitivity phosphoproteomics with dia-PASEF. In the future, these advances must be accompanied by parallel developments in all parts of the workflow to scale phosphoproteomics successfully.

## Acknowledgments

This work was partially supported by the Federal Ministry of Education and Research and the Thuringian Ministry for Economic Affairs, Science and a Digital Society through the Joint Federal Government-Länder Tenure-Track Programme and by the Free state of Thuringia and the European Union via the ‘Innovationszentrum für Thüringer Medizin technik-Lösungen’ (ThIMEDOP; #2018 IZN 002). We thank our colleagues at the Jena University Hospital for fruitful discussions; in particular C. Tschernjawski for technical support.

## Code availability

Custom data analysis and visualization scripts can be accessed via https://github.com/MeierLab.

---

**Supplementary Figure 1.**
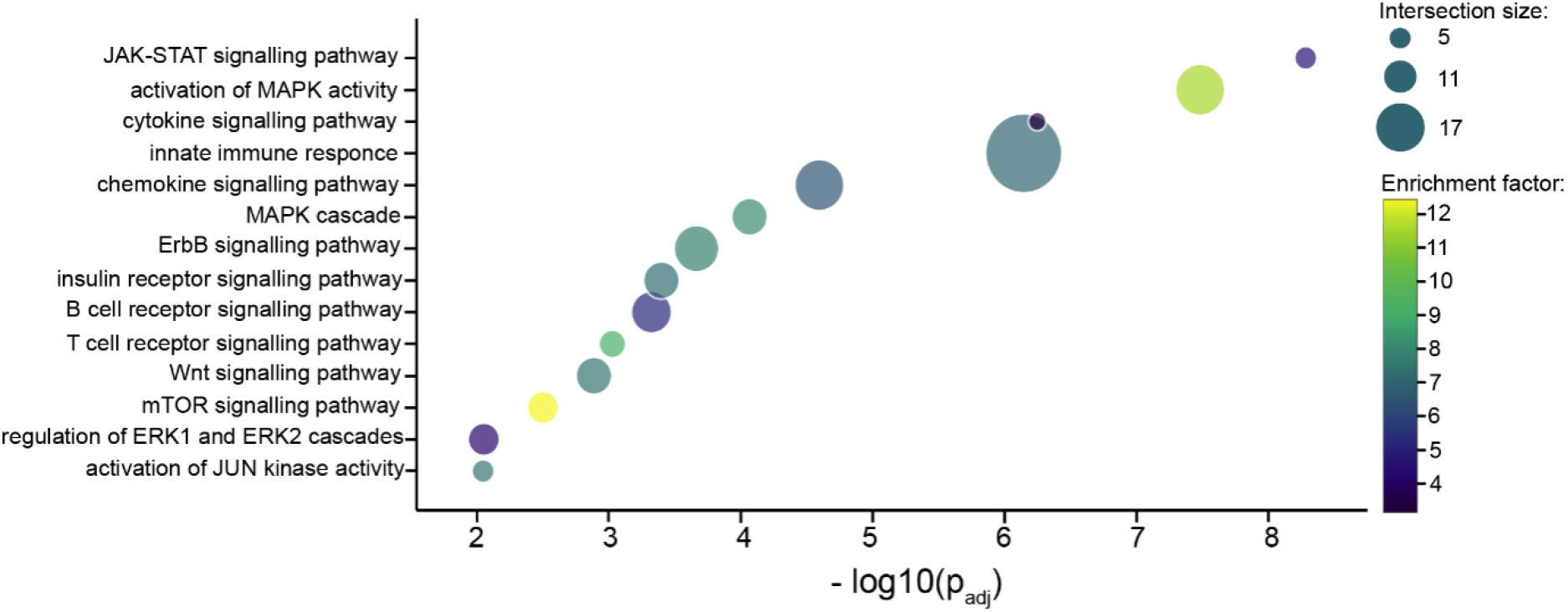
Gene ontology enrichment analysis by Fisher’s exact test for phosphorylated proteins identified in the 7 min gradient.

**Supplementary Figure 2.**
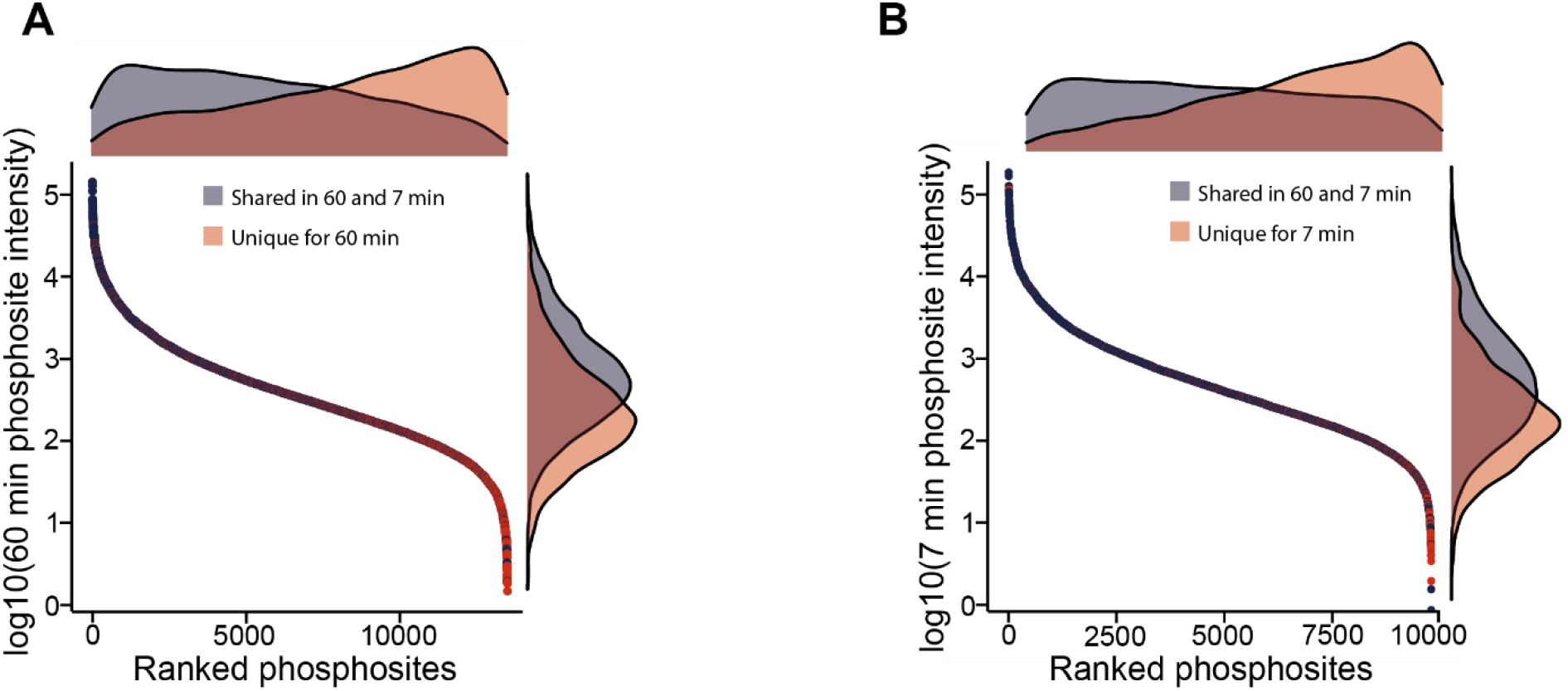
**A** Phosphosite rank plot from highest to lowest abundant sites shared between 60 and 7 min, and unique for 60 min. Kernel densities estimate phosphosite intensities and ranked phosphosites are project on the respective axes. **B**Same as A,but highlighting unique identifications for the 7 min gradient.

**Supplementary Figure 3.**
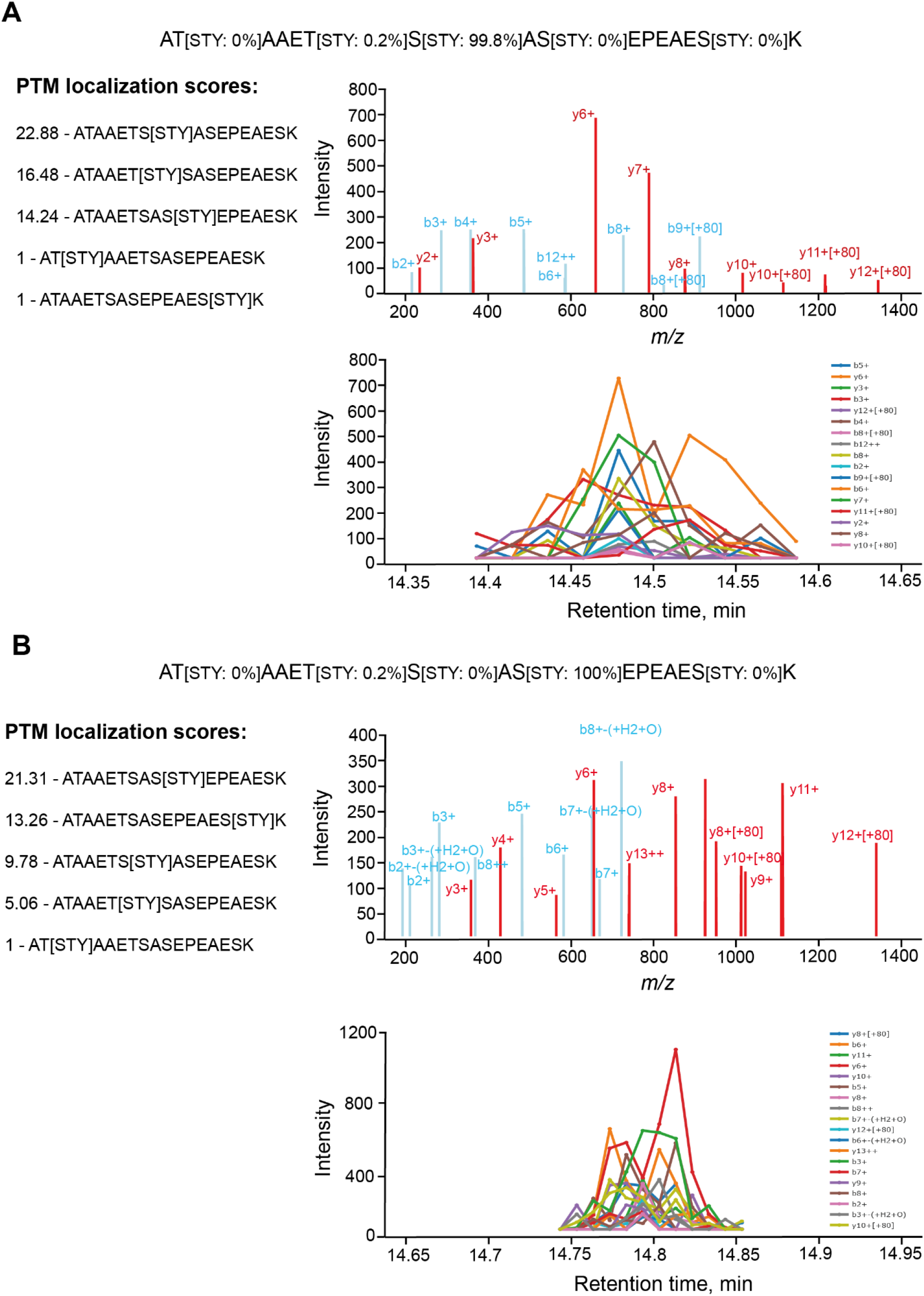
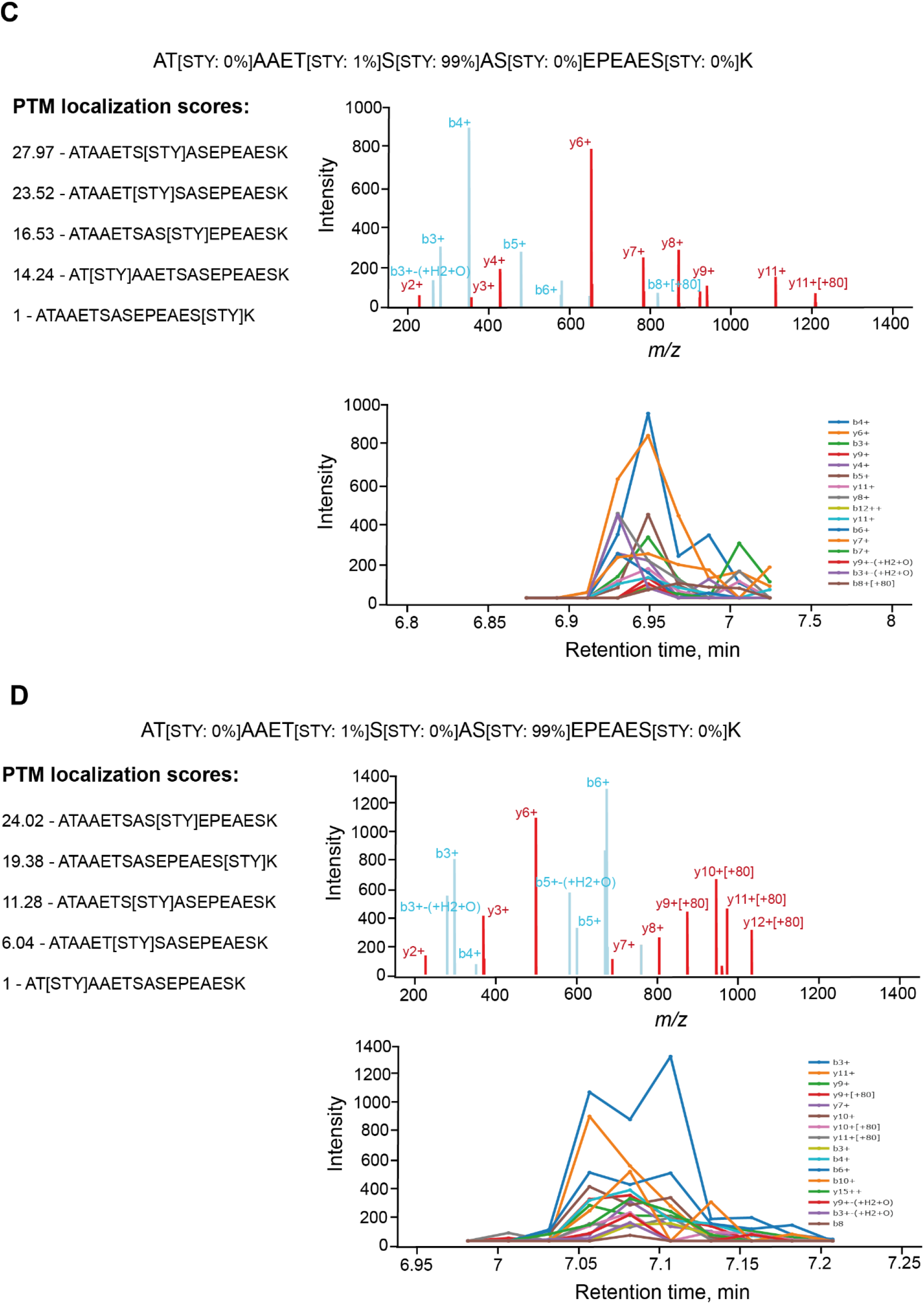
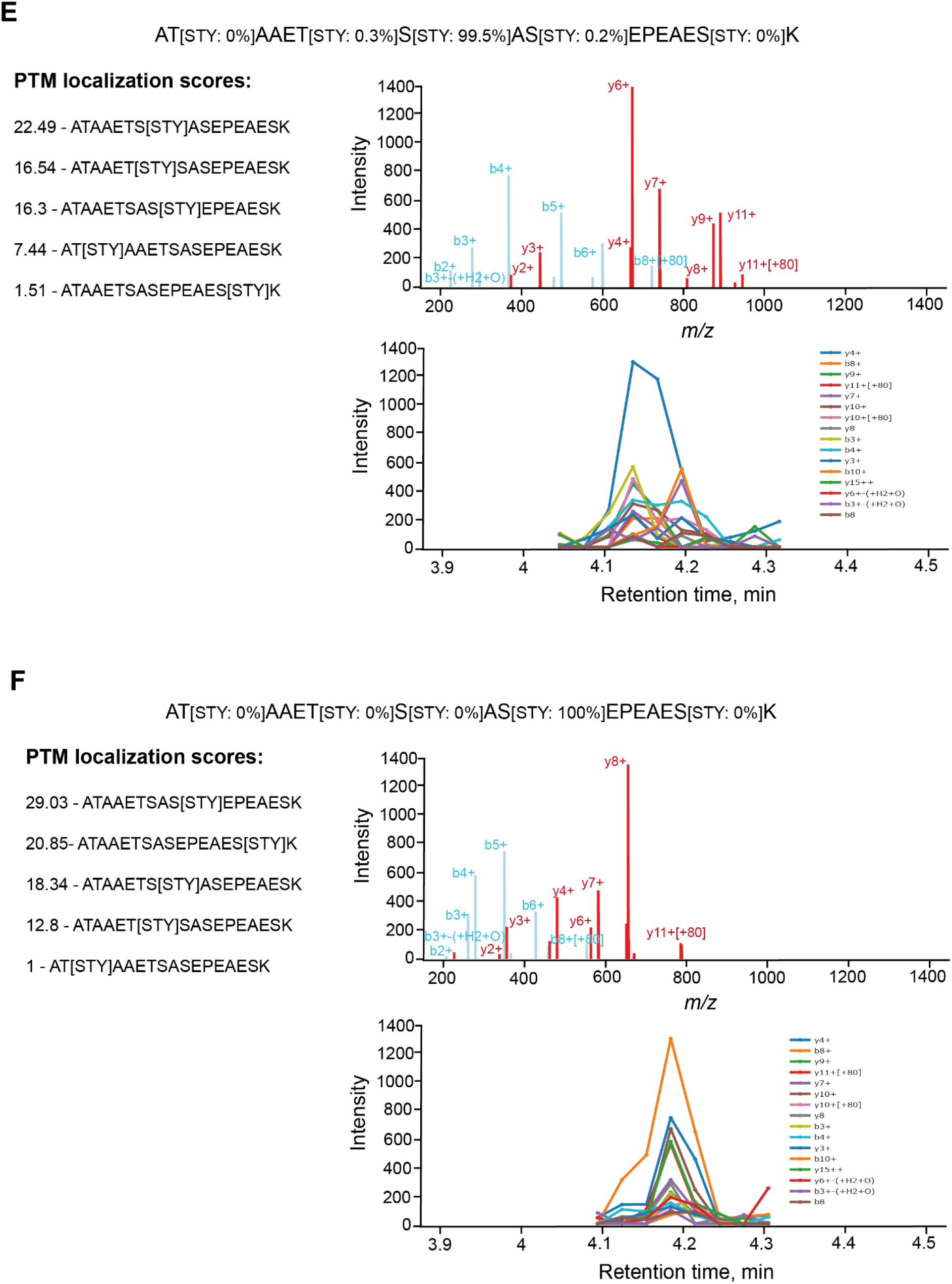
Spectronaut PTM localization plots for the selected phosphopeptides ATAAETpSASEPEAESK and ATAAETSApSEPEAESK identified in 60 min (**A-B**), 21 min (**C-D**) and 10 min (**E-F**) gradients.

